# fMROI: a simple and adaptable toolbox for easy region-of-interest creation

**DOI:** 10.1101/2024.03.29.587330

**Authors:** André Peres, Daniela Valério, Igor Vaz, Morteza Mahdiani, Jon Walbrin, Jorge Almeida

## Abstract

This study introduces fMROI, an open-source software designed for creating regions-of-interest (ROIs) and visualizing magnetic resonance imaging data. fMROI offers a user-friendly graphical interface that simplifies the creation of complex ROIs. It is compatible with various operating systems and enables the integration of user-specified algorithms. Comparative analysis against popular neuroimaging software demonstrates the feasibility, applicability, and ease of use of fMROI. Notably, fMROI’s interactive graphical interface with a real-time viewer allows users to identify inconsistencies and design more accurate ROIs, saving significant time by avoiding errors before storing ROIs as NIfTI files. Additionally, fMROI supports automation through command-line accessibility, making it ideal for large-scale analyses. As an open-source platform, fMROI provides a valuable resource for researchers in the neuroimaging community, facilitating efficient ROI creation and streamlining neuroimage analysis.

## 1. INTRODUCTION

Creating regions of interest (ROIs) is a routine task for anyone working with medical images, and the development of new algorithms is crucial for improving the quality of both clinical and experimental analyses (Hossain et al., 2023; Julian et al., 2012; Pamplona et al., 2020; Pandey et al., 2018; Yogananda et al., 2020).

In functional magnetic resonance imaging (fMRI) research, ROI-based analysis enables researchers to address hypotheses about focally specific brain areas, and alleviates problems associated with en masse multiple comparisons that are inherent to whole brain analyses (Poldrack, 2007). ROIs can be used for various analytic purposes, for example, to measure the magnitude of BOLD activation responses to different stimuli, or the temporal covariation of the BOLD signal between ROIs (Basti et al., 2020; Smith et al., 2013; Stephan & Friston, 2009; van den Heuvel & Hulshoff Pol, 2010; Vieira et al., 2021).

ROIs may be defined in multiple ways – for example, using anatomical atlases (e.g., based on established brain structures; (Destrieux et al., 2010; Fischl et al., 2004; Maldjian et al., 2003), functional atlases based on multi-modal patterns (Glasser et al., 2016; Yarkoni et al., 2011), or guided by the localization of functional responses with independent data (e.g. identifying brain areas that exhibit a preference for face stimuli; Julian et al., 2012; Saxe et al., 2006). Crucially, the method of ROI selection may significantly impact results, even with small variations (Li et al., 2010; Mitsis et al., 2008; Sohn et al., 2015; Tong et al., 2016). Therefore, appropriate ROI choice is essential for both effective hypothesis testing and ensuring reproducible findings (Liu, 2011).

Currently, most fMRI analysis software, such as SPM (https://www.fil.ion.ucl.ac.uk/spm – Friston et al. 2007), FSL (https://fsl.fmrib.ox.ac.uk – Jenkinson et al. 2012), AFNI (https://afni.nimh.nih.gov – Cox 1996), and FreeSurfer (http://surfer.nmr.mgh.harvard.edu – Fischl et al. 2004), include ROI creation options. Although these programs are highly reliable, creating complex ROIs beyond spheres, cubes, or atlas-defined regions can be challenging (e.g., ROIs that consider the intricate shapes of brain activation and the distribution of statistical maps), commonly requiring multiple steps or requiring users to implement their own scripts. In addition, many of these software packages exhibit poor graphical user interfaces (GUI), and some require the use of command-line as their only interface, which makes ROI designing more difficult, less intuitive, and time-consuming.

To address this gap, fMROI aims to facilitate the creation of complex and specific ROIs, making this process more precise and intuitive. It offers a robust GUI integrated with an advanced neuroimage viewing tool and specialized ROI creation tools. Users can design and observe real-time changes in results, ensuring a streamlined and efficient ROI creation experience. Creating ROIs in fMROI is as simple as 1. Clicking on the desired position to select the center of an ROI or entering these coordinates directly in the navigator; 2. Selecting the ROI type (spherical, cubic, region-growing, etc.) and parameters (radius, number of voxels, etc.); 3. Clicking a button to generate the ROI. Combining ROIs or calculating conjunction between them is also very simple in the logical operations tab.

Furthermore, fMROI was designed as a collaborative development platform that caters to the needs of both experienced programmers interested in GUI development and users who create specific tools for their analysis. Its unique architecture consists of three independent modules: 1. The graphical interface, that comprises the viewer, menus, and graphical controls; 2. The ROI methods structure that houses the ROI creation algorithms, and; 3. The tools module that enables extra functionality in fMROI beyond ROI creation, like automated multi-slice capture and image mosaic generation. This modular design allows for seamless updates to the interface while ensuring the smooth functioning of the ROI algorithms.

Another feature that sets fMROI apart from alternative tools is the capacity to effortlessly incorporate new ROI creation algorithms, making it a user-friendly and intuitive platform for importing custom routines. This feature is particularly valuable for biomedicine and neuroscience professionals who, despite varying programming expertise, can easily integrate their analysis code, unleashing the full potential of fMROI as a powerful ally in their research pursuits.

Thus, here, we present fMROI, a new freeware MATLAB-based toolbox for ROI creation that is: 1. Intuitive and easy to use (multiple ROIs can be created with just a few clicks); 2. Versatile (allows for complex and specific ROI creation) and; 3. Adaptive (allows easy integration of specific user-created ROI algorithms). fMROI was tested with simulated and real data and results were compared against other well-established software. In the subsequent sections, we provide a brief overview of existing ROI packages, and an in-depth examination of the fMROI toolbox, presenting its key features, use cases, and extensive validations. For code access, visit https://github.com/peresasc/fmroi, and for a comprehensive user guide, please follow this link: https://fmroi.readthedocs.io.

## 2. OVERVIEW OF EXISTING ROI SOFTWARE

In this section, we provide a concise overview of prominent software packages commonly utilized by the fMRI community for creating ROIs. We specifically focus on four widely recognized software packages, highlighting their key features and significance in the field.

***SPM:*** SPM (Statistical Parametric Mapping – https://www.fil.ion.ucl.ac.uk/spm – Friston et al. 2007) is a widely-used Matlab package for fMRI analysis. Notably, SPM offers the Image Calculator (Imcalc – http://tools.robjellis.net) function, enabling algebraic manipulations among images (e.g., mean or sum images) and binary masking with specific thresholds. Furthermore, SPM benefits from numerous extensions, including MarsBar (https://marsbar-toolbox.github.io – Brett et al. 2002) a popular GUI toolbox. MarsBar facilitates the creation of spherical and rectangular ROIs, as well as their combination and flipping. However, it requires explicit specification of ROI coordinates, lacking the convenience of specifying coordinates by simply clicking on an image. Additionally, MarsBar does not incorporate an image viewer and instead relies on SPM or other external programs for visualization. As a consequence, the process of inspecting and adjusting ROI locations can be time-consuming.

***AFNI*:** AFNI (Analysis of Functional NeuroImages – https://afni.nimh.nih.gov – Cox 1996) is a powerful software available for MacOS and Linux, renowned for its sophisticated GUI and integrated viewer for fMRI analysis. While it offers various algorithms for ROI creation (e.g., spherical, cubic, threshold, clustering, and drawing masks), it’s worth noting that apart from the Draw ROI plugin, all other algorithms are command-line-based. AFNI provides a versatile ROI drawing facility, equipped with multiple drawing tools and the capability to create multiple ROIs and save them as a single labeled image.

***FSL***: FSL (FMRIB Software Library – https://fsl.fmrib.ox.ac.uk – Jenkinson et al. 2012) is a widely utilized package for fMRI analysis, compatible with Linux and MacOS operating systems. FSL allows for ROI creation through its GUI, FSLeyes, as well as via the command line using the fslmaths package. It supports basic ROI types such as spherical, cubic, and threshold masks and includes a functional ROI drawing assistant. On a positive note, fslmaths provides tools for combining ROIs, allowing for further analysis and customization.

***MRIcroGL*:** In contrast to the other software mentioned, MRIcroGL (https://www.nitrc.org/projects/mricrogl – Rorden and Brett 2000) is primarily designed as a brain image viewer rather than an fMRI analysis toolbox. It is compatible with Windows, Linux, and MacOS, offering a user-friendly interface that incorporates a comprehensive set of ROI creation tools. Users can draw ROIs manually on background images, and generate simple ROIs such as spherical and threshold masks. MRIcroGL provides various transformation options, including region dilation and erosion, as well as the capability to combine multiple ROIs, demonstrating its versatility beyond simple image visualization.

Importantly, in contrast to the aforementioned software, fMROI is primarily designed to create ROIs. Its core objective is to integrate the best features of existing software while introducing innovative ROI creation algorithms. Notably, fMROI offers an interactive graphical interface with a real-time viewer, enabling users to refine ROIs efficiently and identify discrepancies before saving them as NIfTI files, enhancing the precision and usability of any neuroimaging analysis.

## 3. METHODS

### 3.1 Dependencies

fMROI has an obvious dependence on Matlab (The MathWorks Inc., Natick, MA – USA), as it was developed within this software. It was systematically tested on Matlab R2023a and R2023b, and maintains backward compatibility, supporting versions as early as R2018a across various operating systems, including Linux (tested on Ubuntu 22.04), MacOS (14.1.1), and Windows 10. fMROI requires the installation of Statistical Parametric Mapping toolbox (SPM12, and is potentially compatible with earlier versions) and Image Processing toolbox (Version 11.5 – The MathWorks Inc., Natck, MA – USA). fMROI also requires the following packages that are already included in its installation: DIRWALK – Walk the directory tree (Prilepin, 2022); findjobj – find java handles of Matlab graphic objects (Altman, 2022); uigetfile_n_dir: select multiple files and directories (Tiago, 2022); get_arg_names (Eaton, 2019).

### 3.2 Implementation

The implementation of fMROI was carefully designed to prioritize user-friendliness and facilitate the addition of new features. To achieve this, a clear separation was established among three main modules: the main GUI implementation, ROI creation algorithms, and extra tools. This modular architecture enables collaborators to introduce new features simply by adding files to specific folders, without the need to understand the inner workings of other parts of the fMROI code.

The functions within fMROI are organized into three distinct categories: GUI creation, Callbacks, and Methods. GUI creation functions are responsible for building graphical interfaces and defining the placement and appearance of objects, such as panels, buttons, and sliders. Callback functions handle user interactions with these objects, executing actions in response to events like button clicks. All other functions not directly related to GUI creation or called through GUI interaction are classified as method functions. To maintain organization, functions are stored in separate folders based on their category (gui, callback, and method folders), as depicted in figure 1A.

**Figure 1.**
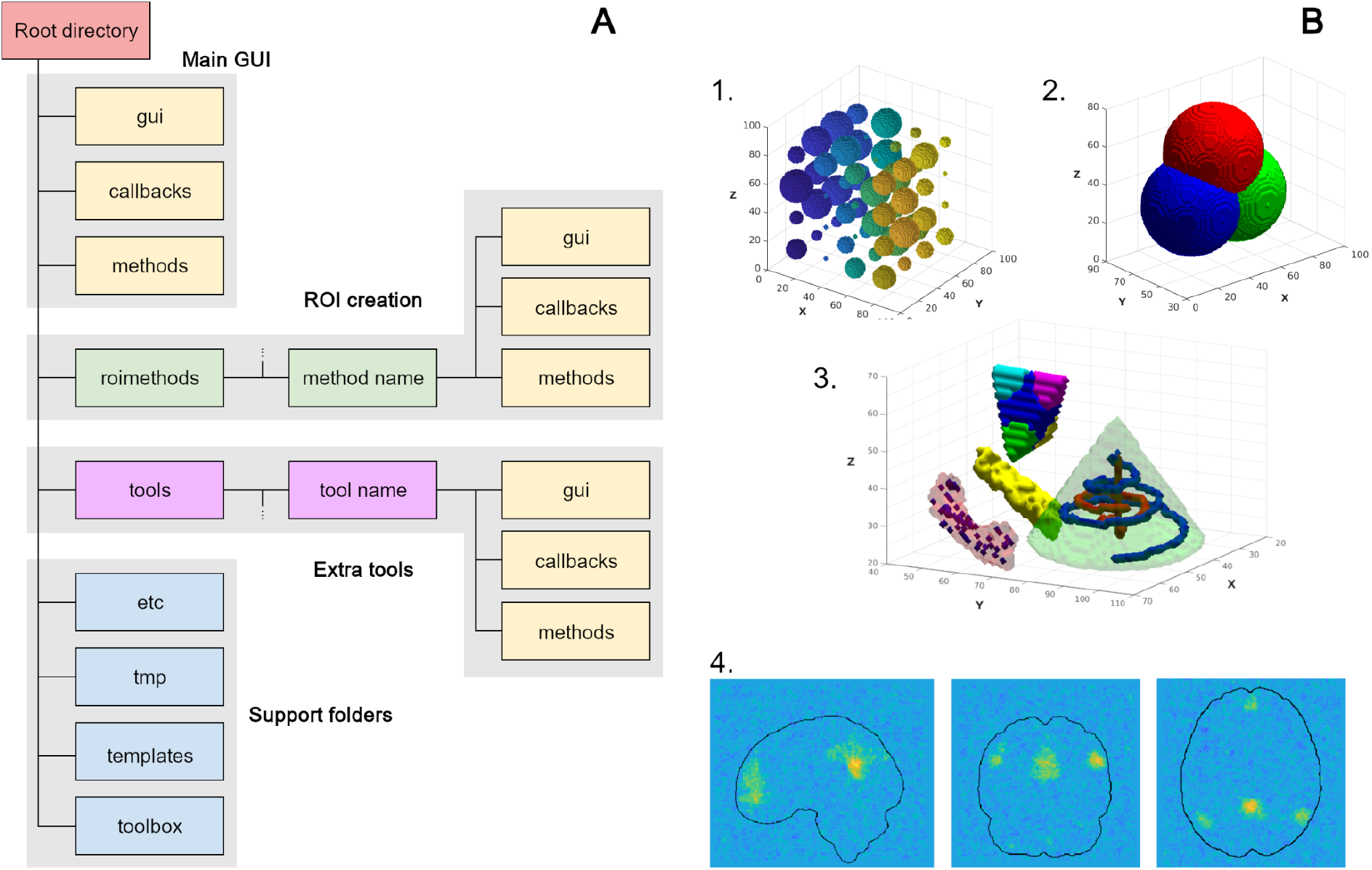
Schema of fMROI file structure and synthetic data used to test the ROI creation algorithms. **A –** fMROI file structure where each box represents a folder. The box in red represents the fMROI root directory, the green box represents the folder that houses the algorithms for creating ROIs, and the boxes in magenta are the extra tools structure. The three sets of yellow boxes represent the directory structure of each of the independent modules, and the blue boxes represent the support structure for the fMROI operation. Each gray shadowed area represents one independent module (main interface, ROI methods, extra tools, and support folders). **B –** Synthetic and hybrid images used to test the ROI creation algorithms.The images from B1 to B3 are three-dimensional reconstructions of the images, while B4 represents the three orthogonal slices. 1. Set of 64 spheres uniformly distributed from the position [13, 13, 13] to [79, 79, 79]. The spheres received a unique integer value ranging from 1 to 64, represented by the colormap from blue (lower values) to yellow (higher values); 2. Set of three images, each containing a sphere, that is partially overlapped. The spheres are centered at the corners of an equilateral triangle and have radius equal to the triangle altitude; 3. Synthetic image composed of several complex shapes that are not necessarily biologically plausible (sharp corners, straight lines, spirals, etc); 4. Hybrid image composed by the Neurosynth DMN association test with the addition of Gaussian noise of approximately 5dB. The outline of the brain is purely illustrative.

This organizational structure is applied across all three modules, each of which is described in detail below:

*Main GUI:* The main interface of fMROI is built on a single frame (Matlab figure), utilizing only uicontrols to ensure compatibility with different versions of Matlab. Builders or graphical assistants such as App Designer or GUIDE were deliberately avoided to maintain code clarity and cleanliness. The main GUI consists of a control panel that lists loaded images and provides controls for adjusting image appearance (e.g., threshold and opacity sliders), three tabs for ROI creation and editing, four axes for displaying planar slices and three-dimensional reconstructions, and a main menu at the top of the frame.

*ROI creation:* The ROI creation functions, located in the “roimethods” folder, are dedicated solely to the creation of ROIs. Default ROI creation algorithms consist of a function triplet comprising the method itself, that carries all the instructions to create the ROIs, the GUI constructor, responsible for generating the graphical interface, and the GUI caller, which integrates the method to the GUI. Only the method function is mandatory and can be invoked either from the interface (via caller function) or directly from the command line, enabling the creation of scripts for batch processing. In the absence of the GUI builder pair (i.e., GUI and Caller functions), fMROI utilizes a generic GUI builder (autogen_gui.m) to generate the graphical interface. For a more detailed explanation of the autogen_gui.m function, refer to the supplementary methods.

*Extra tools:* The extra tools encompass a collection of functions that are not essential for the core functionality of fMROI but can be highly useful in various tasks (located in the “Tools” menu). By default, fMROI 1.0.x includes a screenshot assistant that allows the automated capture of multiple slices and the generation of image mosaics. fMROI also has an import assistant (located in the “Config” menu) to facilitate the addition of new tools. The extra-tools files are stored in the “tools” folder within the fMROI root directory.

Four additional folders – “etc”, “templates”, “tmp”, and “toolbox” – provide essential support for the operation of fMROI. The “etc” folder contains default settings and necessary files for the main interface. The “templates” folder houses template images that can be loaded through the “File > Load Templates” menu. The “tmp” folder is utilized by fMROI for reading and writing temporary files, such as under-construction ROIs. Finally, the “toolbox” folder stores third-party toolboxes used by fMROI (see figure 1A).

### 3.4 Input and output data

The input and output data in fMROI are handled using 3D NIfTI-files. If a 4D NIfTI file is selected, fMROI will only display the first 3D volume from the array. All image loading and saving operations are performed using SPM functions, leveraging the same metafile data structure as SPM (refer to the ‘st’ variable in the spm_orthviews.m function for more details). There are three methods available for loading images in fMROI. Users can choose from preinstalled templates, select an external NIfTI file, or read ROI NIfTI files. In the case of ROI NIfTI files, it is also possible to load a color lookup table (LUT) file containing additional information.

Additionally, fMROI provides the flexibility to import new templates, delete existing templates, or restore the default template set. By default, fMROI 1.0.x includes templates from FSL (FMRIB58_FA-skeleton, MNI152lin_T1, and Talairach – Lancaster et al. 2007), FreeSurfer (aparc+aseg, aparc.a2009s+aseg, and T1 – Fischl et al. 2004; 2002; Destrieux et al. 2010), Neurosynth (images used in this study – default_mode_association-test and faces_association-test – Yarkoni et al. 2011), TemplateFlow (tpl-MNI152NLin2009cAsym and tpl-MNI152NLin6Asym – Ciric et al. 2022), and the synthetic data used in this study (64-spheres, three-sphere set, and complex-shapes images; see figure 1B).

When it comes to saving ROIs, fMROI offers two options. The first is to save ROIs as binary masks, where each individual ROI is stored as a separate NIfTI file. The second is to save ROIs as an indices mask, where all created ROIs are saved within the same NIfTI file, each with a different value (index) assigned to them. It is important to note that the index value must be a positive integer. When selecting the indices mask option, fMROI also saves an complementary LUT file that attributes labels and colors to each ROI, enhancing visualization in the viewer. Regardless of the chosen mode, fMROI saves an information table for each ROI, containing essential details such as the index, label, number of voxels, and mass center coordinates. Additionally, it computes and includes statistical measures – median, mean, and standard deviation – derived from the intensity values of voxels within the source image masked by each ROI. These metrics provide valuable insights into the distribution and characteristics of the underlying neural activity within delineated regions.

### 3.5 ROI algorithms

By default, fMROI 1.0.x has seven algorithms for creating ROIs and one for combining ROIs:

1. *Spheremask* creates a spherical mask filled with non-zero unsigned integer (index) and surrounded by zeros. The center of the spherical ROI is defined by a three-dimensional vector, whereas in the graphical interface, it is represented by the current position of the cursor (curpos). The algorithm allows for the definition of the ROI size by either its radius or volume (number of voxels).
2. *Cubicmask* algorithm works similarly to spheremask but produces a cubic mask. Instead of defining the ROI size by the radius or volume, Cubicmask defines it by the edge or volume.
3. *Maxkmask* searches for the k highest-intensity voxels of the input image constrained in a region defined by a mask (premask). The premask can be a sphere defined by entering its center and radius or a loaded ROI image. If the image selected as a premask is not binary, it will be binarized automatically by the img2mask function.
4. *Regiongrowingmask* is a region-growing algorithm that groups neighboring voxels iteratively according to a rule. This algorithm has three rules for growing: ascending, where it searches for the highest-intensity neighbors; descending where it searches for the lowest-intensity neighbors; and similarity to the seed where it searches for the most-similar voxels (absolute difference) to the seed (the first selected voxel, usually the cursor position). Additionally, there are other two rules for stopping growth: defining a maximum number of voxels to compose the ROI or defining the maximum intensity difference between the seed and its neighbors.
5. *Img2mask* creates a mask determined by the minimum (minT) and maximum (maxT) intensity thresholds (e.g., it generates a mask for brain regions exhibiting activity with a p-value < 0.05). If minT is lower than maxT, img2mask creates a mask with those voxels that have intensity higher than minT and lower than maxT. Otherwise, if the minT is higher than maxT, it creates a mask with those voxels that have intensity higher than minT or lower than maxT.
6. *Contiguousclustering* groups first-neighbors voxels (six-neighbors – voxels that touch each face of the selected voxel), defined by minT and maxT in the same way as in img2mask. Every voxel grouped in a cluster is assigned an index (non-zero unsigned integer) and each cluster is considered an independent ROI. Furthermore, it is possible to eliminate scattered voxels by removing clusters that have fewer elements than a given threshold.
7. *Drawingmask* is a tool for generating ROIs from freehand drawing. ROIs are created layer by layer on planar images and at the end of the procedure, they are stacked into a volumetric ROI. This tool does not have a standalone version, so it only works alongside the fMROI graphical interface.
8. *Logic-chain* performs consecutive logical operations (conjuction, disjunction, and negation) on provided data arrays or NIfTI files. The function computes logical operations iteratively, producing a resultant binary matrix with the same dimensions as the input matrices or NIfTI data arrays. Input data can consist of loaded ROIs and images, which are automatically binarized by the img2mask function, or under-construction ROIs.

### 3.6 Importing new ROI algorithms

An important feature of fMROI is the capacity to import new ROI algorithms because it allows easy incorporation of scripts that previously ran without GUI and stimulates the development of new features. After importing a new method, the *updatepopuproitype* function checks the name of the subfolders within the *roimethods* directory and subsequently incorporates them into the ROI type pop-up menu. Importantly, the method subfolders must have exactly the same name as the main method file. Next, the ROI type pop-up callback function checks if the method includes a GUI creator function. In cases where the GUI creator function is not present, it launches the generic GUI builder (*autogen_gui*) function, which autonomously generates a GUI (for an in-depth explanation of the autogen_gui.m function, please refer to the supplementary methods section).

Alternatively, users have the option to import functions without relying on the fMROI GUI. In such instances, users can easily copy the folder containing the method functions that they wish to import directly into the *roimethods* directory. While it’s not mandatory, to maintain consistency with fMROI code organization, we recommend structuring the method functions as follows: place the method functions within a folder [method name]/methods, place the GUI creation and caller functions in [method name]/gui, and the callback functions in [method name]/callback.

### 3.7 Testing with real and simulated data

In this section, we present the extensive testing performed on fMROI, encompassing both quantitative and qualitative analyses to thoroughly evaluate its functionalities and performance. The quantitative analyses primarily focused on validating the accuracy and precision of the ROI creation algorithms, while the qualitative analyses served to demonstrate the diverse features and capabilities of fMROI.

#### 3.7.1 Testing Dataset

To test each ROI creation algorithm, we prepared six images: an MNI structural T1-weighted from FreeSurfer (256×256×256 voxels with isovoxel of 1 mm^3^); two fMRI maps collected from Neurosynth meta-analysis database (https://neurosynth.org – Yarkoni et al. 2011), both 91 x 109 x 91 voxels with isovoxel of 2 x 2 x 2 mm^3^); and three synthetic data images (91 x 109 x 91 voxels with isovoxel of 2 x 2 x 2 mm^3^; see figure 1B).

To obtain the Neurosynth data, we input the search term “default mode” which returned 777 studies and 26256 activations (here referred to as DMN image), and the term “faces” which returned 864 studies and 31842 activations (faces image). We then downloaded the association test images.

To generate the synthetic data we created: 1. *Three-sphere set images:* a set of three binary NIfTI files, each containing a sphere centered at one of the corners of an equilateral triangle; 2. *64-spheres image:* a single NIfTI file with 64 spheres of different sizes uniformly distributed; and 3. *complex-shapes image:* a sigle NIfTI file that combines geometric shapes (tetrahedron, cone, and spirals) with left and right hippocampi sourced from aparc+aseg atlas (Fischl et al., 2002). Specifically, the left hippocampus is uniformly filled with ones, while the right hippocampus contains randomized values ranging between 3 and 4. In the tetrahedron, the central region uniformly holds a value of 5.0, whereas its four corners exhibit values of 5.2, 5.4, 5.6, and 5.8, respectively. The cone has random values from 9 to 10, with two spirals connected by a straight line inside of it. The external spiral has values ranging from 11 to 12 from the cone’s tip to the base, the internal spiral has values ranging from 7.17 to 8, increasing from cone’s base to tip, and the connecting line with values increasing from 7.0 to 7.16 from cone’s tip to base. All synthetic images were created on a 91 x 109 x 91 matrix with a voxel size of 2 x 2 x 2 mm^3^ (see the supplementary methods for a detailed explanation of the synthetic data creation).

#### 3.7.2 Validation tests – Quantitative analysis

To test the ROI creation algorithms, we compared the generated outputs with theoretical values or calculated the F1-score by comparing the outputs with pre-selected templates.

Specifically, for spherical and cubic ROI analysis, we compared the theoretical area and volume to the measured area and volume (for details about area and volume calculation, check the function roi_areavol at [fMROI root folder]/etc/scripts). Additionally, we assessed the alignment of the ROI center of mass with the specified ROI center position. For the img2mask, maskmask, and drawmask algorithms, the ground truth could be easily determined by applying fundamental mathematical relationships such as greater than (>), less than (<), or equal to (=) to virtually any image. However, to create the ground truth for the contiguouscluster algorithm, we had to generate a synthetic NIfTI image with 64 non-contiguous regions (*64-spheres image*), each with distinct and known values. Similarly, for generating the ground truth for the regiongrowing algorithm, we created a NIfTI image with several shapes (*complex-shapes image*). The values were carefully chosen with the intention of guiding the ROI growth in a specific direction, or to halt growth when encountering regions with values exceeding the seed’s difference threshold. All data and scripts used for the quantitative and qualitative analyses can be accessed in the supplementary data at the Zenodo repository https://doi.org/10.5281/zenodo.10845325 or in the GitHub project https://github.com/peresasc/fmroi_supplementary_data. For details on all the quantitative tests conducted, please refer to the script *[supplementary_data]/scripts/validation/roigen_qc.m*.

##### Spherical and Cubic ROIs test

To evaluate the spherical and cubic mask methods, we generated 100 ROIs (radius/edge varying from 1 to 100 voxels and random centers) for each software (fMROI, SPM, MRIcroGL, FSL, and AFNI). Each ROI was stored in a different NIfTI file using the FreeSurfer T1 image as the template. For each parameter pair (radius/edge and center of mass), we computed the theoretical sphere/cube within a continuous space. Next, we calculated three independent linear models to explain the areas, volumes, and centers of mass of the ROIs created by the algorithms concerning their respective theoretical values. Subsequently, we measured the deviation of the calculated models from the ideal y = x model (i.e., observed value equals theoretical value) using the Normalized Root Mean Squared Deviation (NRMSD – see Table 1).

**Table 1:**
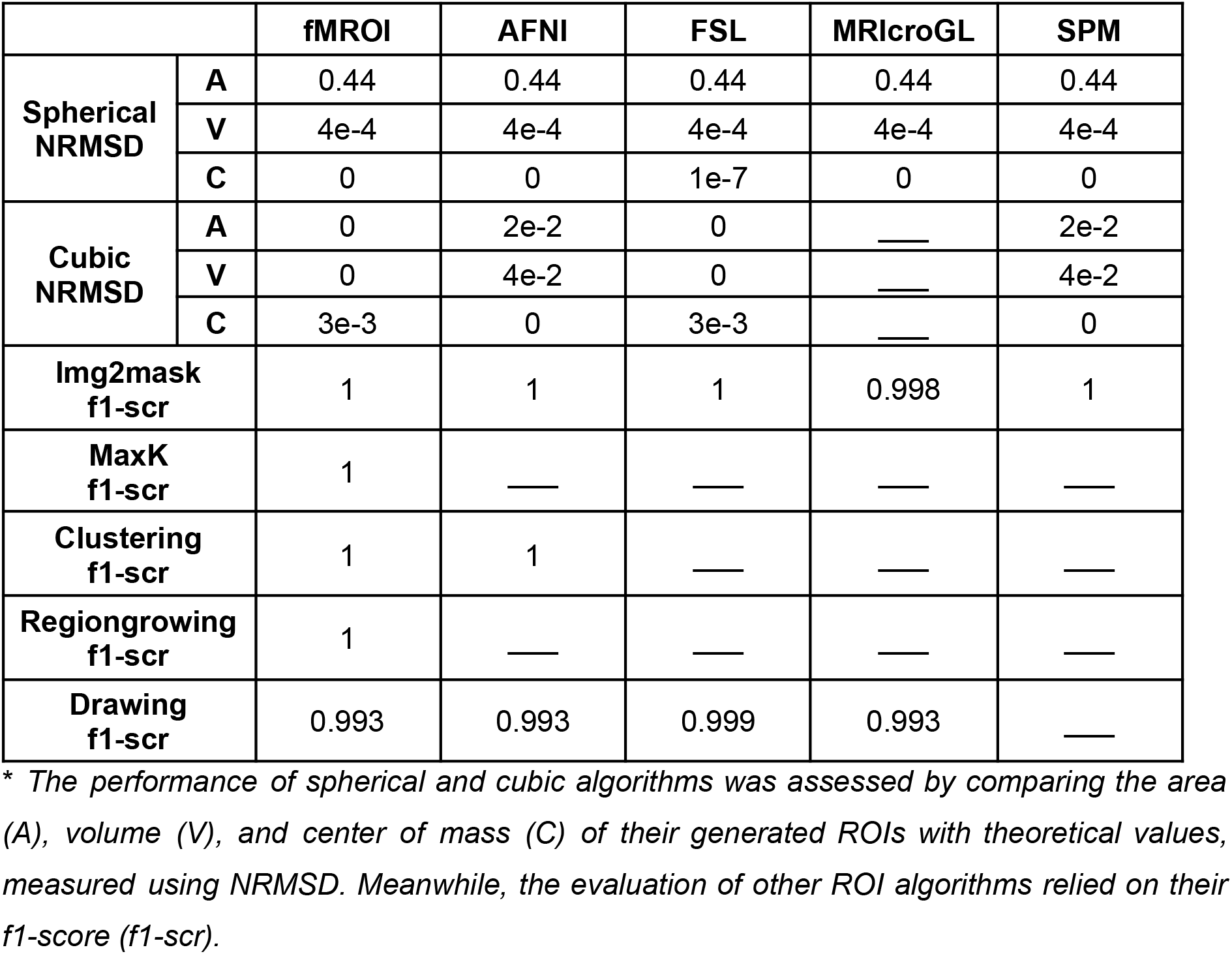
Summary of Quantitative Results Across Tested Software.

##### Image threshold mask

To evaluate the image threshold mask methods (img2mask), we used the *complex-shapes* image (figure 1B3) and DMN image. For the *complex-shapes* data, we set the following pairs of minT and maxT to select each shape separately: {1, 1}; {3, 4}; {5, 5}; {5.2, 5.2}; {5.4, 5.4}; {5.6, 5.6}; {5.8, 5.8}; {7, 8}; {9, 10}; {11, 12}. For the DMN, we generated 100 pairs of minT and maxT with values that could vary randomly from the minimum to the maximum value of the source image. In total, we generated 110 ROIs (10 from complex-shapes data and 100 from DMN) to evaluate the img2mask methods.

##### Cluster ROIs test

To evaluate the cluster methods, we used the *64-spheres* image as template (figure 1B1). We defined three sets of parameters {minT, maxT, mincsz} – {0.1, inf, 1}; {17, 32, 33}; {33, inf, 123}, which generated 100 ROIs in total. Next, we evaluated whether the clustering algorithms were able to create the ROIs and whether they clustered all elements.

##### Maximum k elements test

To evaluate the maxk methods, we added Gaussian noise with an SNR of approximately 5dB to the DMN image and used it as the template (see figure 1B4). We used all 112 regions from the FreeSurfer aparc+aseg atlas as premasks. We constrained the search for the highest k-voxels to the region defined by each premask, and the k-highest voxels were defined randomly in the interval of 1 to the maximum number of elements in the premask.

##### Region growing methods test

To evaluate the region growing methods, we used the complex-shapes image in three different approaches. First, we tested the ascending, descending, and similarity growing strategies, stopping the growth according to a pre-established number of elements (similar to the maxk method but considering only connected voxels). We used the tip of the external spiral as the seed [46, 84, 50] to test ascending and descending algorithms, selecting 20 different ROI sizes for each strategy (31 to 221 voxels in steps of 10 for ascending and 21 to 116 voxels in steps of 5 for descending). To test the similarity strategy, we used both the external spiral tip [46, 84, 50] and the voxel with the lowest value of the spirals’ union bar [46, 84, 49] as seeds. For each condition, we selected 10 ROI sizes (23 to 221 voxels in steps of 22 for the seed [46, 84, 50], and 16 to 115 voxels in steps of 11 for the seed [46, 84, 49]). In total, we generated 60 ROIs.

In the second test, we tested the ascending, descending, and similarity growing strategies when the growth is stopped by the difference with the seed (only voxels with a difference from the seed lower than the threshold are considered to compose the ROI). We drew a random voxel from each of the 10 subsets of the complex-shapes data ({x=1}; {3≤x≥4}; {x=5}; {x=5.2}; {x=5.4}; {x=5.6}; {x=5.8}; {7≤x≥8}; {9≤x≥10}; {11≤x≥12}) and used it as a seed for each one of the three growing strategies (ascending, descending, and similarity), generating 30 ROIs in total. The difference threshold was defined as 0.001 for those subsets where all elements have the same value, and 1 for the other subsets.

Finally, in the third test, we performed the same procedures as in the second test, but we also took into account premasks to restrict growth. The premasks were four spherical ROIs ([58,57,27] r=8; [33,57,27] r=8; [47,61,56] r=9; [46,84,58] r=15) that partially overlapped the four main shapes (left and right hippocampus, tetrahedron, and cone).

##### Drawing ROIs test

To assess the efficacy of the drawing methods, we asked six volunteers with different backgrounds to draw the left hippocampus using the *complex-shapes* image. Subsequently, we compared the resulting ROIs with the corresponding elements of the actual left hippocampus in the complex-shapes image.

#### 3.7.3 Application to real data – Qualitative analysis

For the qualitative analysis, we utilized the Neurosynth faces image. To ensure clear separation among brain activations, we applied a minimum threshold equal to half the maximum value of the image (z-score = 21.46) rounded to the nearest integer (minT = 11). The maximum threshold was set to the maximum value of the image (maxT = 21.46). Using the “Find max” function, we identified the voxel with the highest associated value (MNI [42, –50, –24]; data matrix in LAS coordinates [25, 39, 25]), which served as the seed for the seed-based algorithms (spheremask, cubicmask, and regiongrowingmask).

Specifically, for the spheremask test, we set the radius to 3 voxels, resulting in a total of 123 voxels in the ROI. In the cubicmask test, we selected an edge length of 5 voxels, creating an ROI of 125 voxels. For the regiongrowingmask, we employed the ascending strategy without any constraints, resulting in an ROI of 125 voxels. The maxkmask test generated an ROI with 125 voxels without any premasking. In the contiguousclustering test, we did not impose a minimum cluster size (minimum cluster size = 0). To evaluate the drawingmask, a single user drew the activation cluster containing the voxel with the highest value (MNI [42, –50, –24] – right fusiform gyrus). Lastly, for the image2mask method, no additional parameters were required.

To test the *logic-chain* algorithm, we employed the three-sphere set image (see figure 1B2). Each sphere was considered as a separate ROI: S1, S2, and S3. We performed the following logical operations: Disjunction of S1, S2, and S3 (S1 OR S2 OR S3), resulting in the ROI S1∨S2∨S3; Conjunction of S1 and S2 (S1 AND S2), resulting in the ROI S1∧S2; Conjunction of S1 and S3 (S1 AND S3), resulting in the ROI S1∧S3; Conjunction of S2 and S3 (S2 AND S3), resulting in the ROI S2∧S3; Conjunction of S1, S2, and S3 (S1 AND S2 AND S3), resulting in the ROI S1∧S2∧S3; and Conjunction of the Negation of S2∧S3 with the Disjunction of S1∧S2 and S1∧S3 (S1∧S2 OR S1∧S3 NOT S2∧S3), resulting in the ROI [S1∧S2]∨[S1∧S3]∧¬[S2∧S3].

## 4. RESULTS

fMROI was systematically tested on multiple operating systems (OS – Ubuntu 22.04, Windows 10, and MacOS 14) using Matlab R2023a and R2023b. The stability of fMROI was evaluated on all tested OS and Matlab versions, and it demonstrated stability across the board. Notably, the Linux PCs used in the tests were equipped with the NVIDIA GP107GL – Quadro P1000 graphics card, and to achieve good rendering results in Matlab, it was necessary to replace the pre-installed graphic drivers (nouveau) with NVIDIA’s proprietary drivers (driver version: 515.65.01). Additionally, on MacOS with an ARM processor, we encountered some difficulty installing SPM12 when downloaded directly from the official SPM website. We used a workaround by downloading the Development Version from GitHub (https://github.com/spm/spm) and then removing the SPM .mex files from quarantine.

### 4.1 Interface Overview

Figure 2 illustrates the fMROI interface, highlighting its three control sections: A1. Listing table of loaded images, A2. Control of image visualization, and A3. ROI generation and manipulation section. Figure 2A4 showcases a script that demonstrates a simple function for creating cylindrical ROIs (check this function at [fMROI root folder]/etc/scripts/simplecylidermask.m), emphasizing how fMROI automatically generates a GUI for such scripts. The script includes four input arguments, with “srcvol” and “curpos” being fMROI keywords that receive internal values automatically, while “radius” and “height” are presented in the interface as text boxes for manual input.

**Figure 2.**
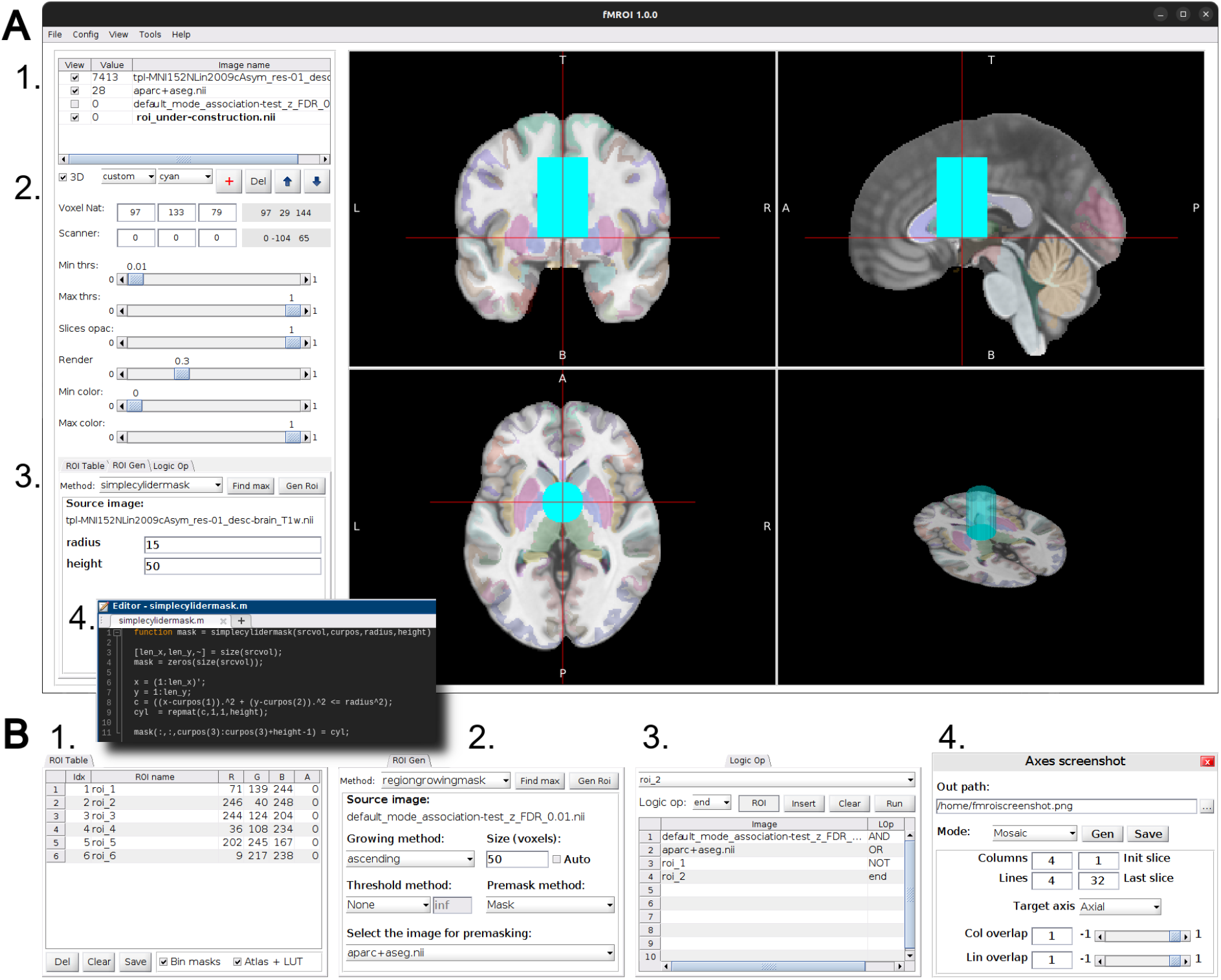
fMROI Graphical User Interface (GUI). **A –** Main interface emphasizing the three control sections and the import of ROI generation scripts. 1. Listing table of loaded images in the order of overlay. In this section it is possible to select images, enable or disable their visualization and visualize the value of the voxel pointed by the cursor; 2. Control of the image visualization aspect. Here it is possible to define colormap, threshold, transparency, cursor position, and stacking order of images. 3. ROI generation and manipulation section. It is divided into three tabs detailed in B (ROI Table, Gen ROI and Logic Op); 4. Example of a simple script that can be easily imported into the fMROI interface. This script generates a cylindrical ROI on an image with the same dimensions as srcvol. The red line highlights that the input variables “radius” and “height” are automatically translated in the GUI as an input text box while the variables “srcvol” and “curpos” receive internal system values as they are fMROI keywords. **B –** Three tabs for generating and manipulating ROIs and axes screenshot interface (Extra tools). 1. *ROI Table* tab lists all ROIs that are under-construction. This tab contains an editable lookup table where it is possible to insert the labels, indices, color and transparency of each ROI. It also contains buttons to delete and save ROIs in NIfTI files; 2. *ROI Gen* tab is the interface to generate new ROIs. It has a drop-down menu to select ROI generation algorithms and a function to find the highest value voxel. For each algorithm there is a GUI for selecting input parameters; 3. *Logic Op* tab is the interface to combine ROIs. Here, it is possible to perform the three logical operations Conjunction (AND), Disjunction (OR), and Negation (NOT). It is also possible to combine under-construction ROIs with loaded images; 4. Axes screenshot assistant is an interface within the ‘Extra tools’ module designed for capturing images displayed in the axes. It offers three distinct modes: mosaic (illustrated here), multiple slices, and single slices.

The bottom part of figure 2 (part B) presents three tabs for creating and manipulating ROIs. The first tab (B1) is similar to section A1 but lists the under-construction ROIs instead of the loaded images. The ROI Table in this tab is editable, allowing manual assignment of ROI indices, names, colors (RGB values), and transparency (alpha). In this example, six ROIs are depicted, and upon completing the ROI creation, users can save them as independent binary NIfTI files by ticking “bin masks” or save them in a single NIfTI file with different indices by ticking “Atlas + LUT” (atlas-like image plus lookup table with ROI Table values).

Figure 2B2 shows the tab for generating ROIs, where the selected algorithm in the Method dropdown menu determines the displayed interface for setting algorithm parameters. It is important to note that this tab includes a “Find Max” button, which finds the voxel with the highest value based on the selected algorithm and its parameters, i.e., the search is limited to the ROI defined by the interface parameters, even if no ROI is created in practice.

The Logic Op tab enables logical operations between ROIs. Figure 2B3 demonstrates a chain of three logical operations: 1. Conjunction (AND) of two loaded images, 2. Disjunction (OR) of the previous operation with an under-construction ROI, and 3. Conjunction of the previous output with the negation (NOT) of another under-construction ROI. Notably, fMROI allows performing logical operations between both loaded images and under-construction ROIs. If the loaded images are not binary, they are automatically binarized using the img2mask algorithm based on the selected MinT and MaxT.

### 4.2 Validation tests – Quantitative analysis

Quantitative analyses reveal exceptional performance for fMROI, consistently achieving maximum scores across most ROI creation algorithms (Table 1). While the other evaluated software (SPM, AFNI, FSL, and MRIcroGL) also demonstrate high performance, fMROI possesses unique functionalities. Notably, it is the only software to offer dynamic ROI projection within both its graphical user interface (GUI) and command-line interface (enabling automation) for all algorithms. Furthermore, fMROI stands alone as the software specifically designed to seamlessly import new ROI creation algorithms. This includes a range of strategies to streamline algorithm incorporation, eliminating the need for users to program the GUI.

The spheremask and cubicmask algorithms were tested by comparing the center of mass, area, and volume of their respective ROIs with theoretical values. All spheremask algorithms (fMROI, SPM, AFNI, FSL, and MRIcroGL) achieved the same results: NRMSD of 0.44 for the surface area, 4e-4 for volume, and 0 for the center of mass. Deviations in area and volume are justified by the discrete space (2×2×2 mm3 resolution image), which introduces corners in the ROI surface, resulting in small increases in the resulting area. Nevertheless, the volume deviation was very small, demonstrating that the theoretical volume is preserved as far as possible. For more detailed information and visual representation of these findings, please refer to the supplementary results section (figure S1).

The cubicmask algorithms (fMROI, SPM, AFNI, and FSL) obtained similar scores, with a slight variation between two groups of algorithms – those optimizing area and volume and those optimizing the center of mass. Due to the discrete nature of digital images, when the number of voxels on the edges is even, the center of mass coordinates are fractions. There are two ways to approach this issue: 1. keeping the selected center and accepting only edges with an odd number of voxels or 2. accepting odd and even numbers of voxels in the edges but resulting in a displacement between the theoretical and selected center.

fMROI and SPM use the second strategy, achieving an NRMSD of 0, 0, and 3e-3 for area, volume, and center of mass, respectively. This means there is a perfect match between theoretical and measured area and volume, but a small displacement between the theoretical and measured center of mass. In contrast, AFNI and FSL use the first strategy, obtaining an NRMSD of 2e-2, 4e-2, and 0 for area, volume, and center of mass, respectively. This means there is a perfect match between the theoretical and measured center of mass, but a small variation between the theoretical and measured area and volume (Table 1). When analyzing only ROIs with edges having odd numbers of voxels, all cubicmask algorithms achieved a perfect match (NRMSD = 0) for all evaluated features (see supplementary results – figures S2 and S3).

Importantly, fMROI algorithms such as img2mask, contiguousclustering, maxkmask, and regiongrowingmask produced an f1-score of 1, indicating that the generated ROIs were identical to the template ROIs used as ground truth (Table 1). For the drawingmask algorithms we obtained an f1-score above of 0.99 for all tested software, indicating that all algorithms have good reproducibility.

### 4.3 Application to real data – Qualitative analysis

In the qualitative analysis, fMROI performance was evaluated using meta-analysis fMRI data related to “faces” (Neurosynth search term: “faces”; association-test FDR = 0.01). The voxel coordinate with maximum activation ([42, –50, –24]) was used as the seed, and a threshold equal to half of the maximum activation (minT=11) was applied. The qualitative results were consistent with the quantitative findings. The *img2mask* and *contiguousclustering* algorithms showed a perfect match between the activation map and the generated ROIs. The *contiguousclustering* algorithm perfectly separated all non-contiguous regions. The ROIs generated by the *spheremask* and *cubicmask* algorithms visually resembled a sphere and a cube, respectively, centered on the seed coordinate. The *maxkmask* and *regiongrowingmask* algorithms produced similar results, with their ROIs covering the activation region containing the highest activation voxel (right fusiform). However, the ROI produced by the *maxkmask* algorithm also covered part of the activations corresponding to the right and left hippocampi, as *maxkmask* can produce sparse ROIs unlike the *regiongrowing* algorithm.

Finally we tested the *logic-chain* algorithm using the three-sphere images set. These spheres are partially overlapped and we performed the following operations: 1. Disjunction of S1, S2, and S3 (figure 3C1); 2. Disjunction of the Conjunctions S1 and S3, S1 and S2, and S2 and S3 (figure 3C2); 3. Conjunction of the Negation of S2∧S3 with the Disjunction of S1∧S2 and S1∧S3 (figure 3C3); and 3. Conjunction of S1, S2, and S3 (figure 3C4);

**Figure 3.**
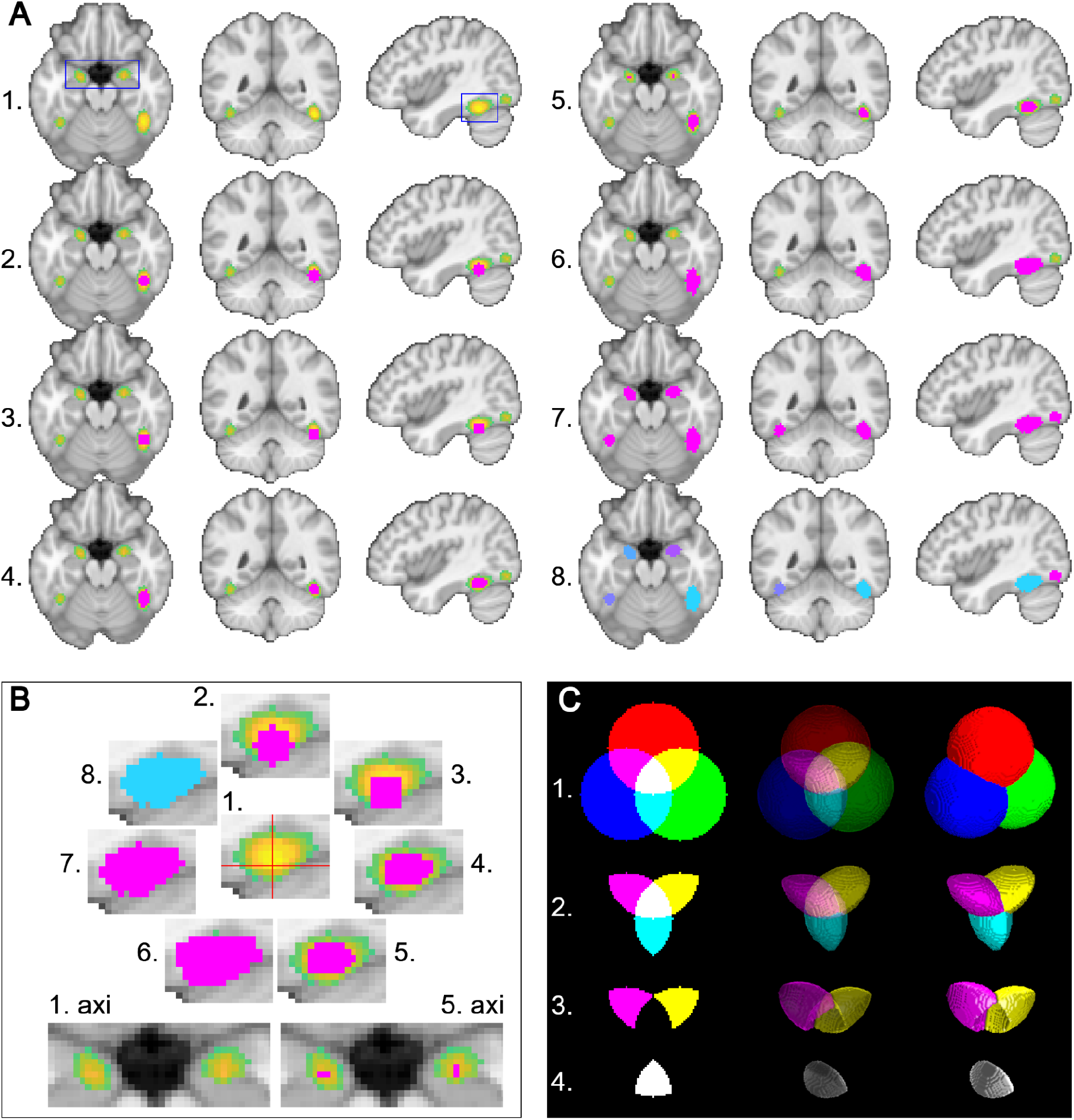
Graphical representation of qualitative results. **A –** Axial, coronal, and sagittal images of the Neurosynth faces association test overlayed by the ROIs generated with the default fMROI algorithms. The slices shown are in neurological representation (left hemisphere is represented in the left side of the image) at MNI coordinates [42,-50,-20], thresholded by z-scores > 11, and presented as: 1. Neurosynth image with no overlay, the rectangles in blue are zoom windows depicted in part B; 2. ROI generated by the *spheremask* algorithm (center = MNI [42,-50,-24]; radius = 3 voxels); 3. ROI generated by *cubicmask* algorithm (center = MNI [42,-50,-24]; edges = 5 voxels); 4. ROI generated by the *regiongrowingmask* algorithm (center = MNI [42,-50,-24]; volume = 125 voxels; ascending); 5. ROI generated by the *maxkmask* algorithm (volume = 125 voxels, no pre-masking); 6. ROI generated by the *drawingmask* algorithm (target = activation cluster at right fusiform gyrus); 7. ROI generated by the *image2mask* algorithm (minT = 11; maxT = 21.46); 8. ROIs generated by the contiguous clustering algorithm (minimum cluster size = 0). **B –** Enlarged view of the regions represented by the blue rectangles on A1 axial and sagittal slices. The upper part, numbered from 1. to 8., represents the clippings respectively of the sagittal images from A1 to A8. All clippings are in the same position and evince the activation cluster in the right fusiform. The lower part shows two clippings, 1. axi and 5. axi. These images respectively represent clippings of the A1. axial slice and A5 axial slice, and emphasize the activation clusters in the right and left hippocampus. **C –** Graphical representation of the results of the *logic-chain* algorithm analysis. The lines (1. to 4.) are the ROIs resulting from the logical operations depicted from left to right in the coronal planar image (MNI [0,0,0]), volumetric rendering with transparency, and volumetric rendering without transparency. 1. Disjunction of the three spherical ROIs used in the *logic-chain* test, sphere 1 (S1) is represented in red and centered at MNI [0,0,32], sphere 2 (S2) in green at [28,0,-16], and sphere 3 (S3) in blue at [-28,0,-16]. 2. Disjunction of the three Conjunctions [S1∧S2]∨[S1∧S3]∨[S2∧S3] 3. Conjunction of the Negation of S2∧S3 with the Disjunction of S1∧S2 and S1∧S3, [S1∧S2]∨[S1∧S3]∧¬[S2∧S3]. 4. Conjunction of S1, S2, and S3, S1∧S2∧S3.

In figure 3C1, we show the Disjunction of the three spherical ROIs – colors were merely used to indicate which sphere each voxel belongs to since the *logic-chain* outputs are binary masks. Thus, red voxels belong only to ROI S1, green to S2, blue to S3, yellow to S1 and S2, magenta to S1 and S3, cyan to S2 and S3, and white voxels belong to the three spherical ROIs. The Conjunction of each sphere pair resulted in three convex solids (similar to a circular double convex lens) and the Disjunction of them resulted in a three-petal solid depicted in figure 3C2. Figure 3C3 shows a two-petal shape with a concave that resulted from the Conjunction of the Negation of the Conjunction of S2 and S3, with the Disjunction of the Conjunctions S1 with S2 and S1 with S3. The Conjunction of the three spherical ROIs resulted in an oblong shape with three sharp edges and two corners depicted in figure 3C4. We chose these three spherical ROIs as the basis for logical operations because their results are predictable and easily evaluated by visual inspection. Thus, all the outputs of the *logic-chain* algorithm were as expected.

In summary, the extensive validation tests demonstrate that fMROI excels in capturing ground truth data across each of the implemented ROI creation algorithms, showcasing its reliability and versatility in analyzing fMRI images. The quantitative analyses revealed exceptional performance across various metrics, with fMROI consistently achieving maximum scores for most ROI creation algorithms. Additionally, qualitative assessments further underscored the software’s efficacy, with findings mirroring quantitative results. Overall, these results affirm fMROI’s robustness and effectiveness in facilitating precise and comprehensive ROI generation for fMRI analysis.

## 5. DISCUSSION

We developed, tested, and validated fMROI – a user-friendly free software for creating ROIs and visualizing brain images. Exhaustive validation tests demonstrate that fMROI produces flawless results in both simulated and real data across all implemented ROI creation algorithms, demonstrating that it is a versatile, flexible, and reliable tool for fMRI analysis. As emphasized by Cohen (2017), while other languages like Python and C++ offer similar or even superior capabilities for some tasks, MATLAB’s established dominance in cognitive neuroscience research fosters a rich ecosystem of powerful and well-supported toolboxes specifically designed for fMRI analysis (Friston et al., 2007; Nili et al., 2014; Oosterhof et al., 2016; Whitfield-Gabrieli & Nieto-Castanon, 2012).

One common requirement of scientific programming is that developers possess a deep understanding of the application, often leading researchers to create their own routines. However, the vast majority of neuroscience researchers lack formal programming education, and as such, knowledge is variable across the field (Hannay et al., 2009). This is where fMROI stands out. By providing a platform where users can easily integrate their own algorithms directly into the interface, fMROI addresses two key issues: firstly, it eliminates the need for researchers to have GUI development knowledge. This makes fMROI accessible to a wider range of researchers with varying programming expertise. Secondly, it encourages researchers to share their code, which can lead to further improvements in the tool (Barnes, 2010).

Hence, fMROI was intentionally developed as a collaborative platform, with a meticulously designed data structure that shields the interface against disruptions stemming from updates to newly embedded algorithms. This protective framework extends not only to the integration of novel ROI creation algorithms but also to the incorporation of template images and extra tools that add new functionalities to the software.

Moreover, as emphasized by Wilson et al. (2014), software should be regarded as an experimental apparatus, subject to scrutiny and testing, akin to any physical instrument. The public release of source code enables the broader community to engage in collaborative evaluation alongside the developers. This transparency benefits not only the research community, ensuring clarity in applied operations and reproducibility of the results, but also the developers, who can iteratively enhance their software through community feedback (Eglen et al., 2017; Gleeson et al., 2017). To make it possible, we have made all the data and code used for fMROI’s testing readily accessible (https://zenodo.org/doi/10.5281/zenodo.10845324).

In our tests, fMROI demonstrated flawless performance in both quantitative and qualitative analysis when compared to well-established neuroimaging software such as SPM, AFNI, FSL, and MRIcroGL. The results consistently showed comparable scores across these platforms. Moreover, fMROI is a dedicated ROI creation tool that implements a broader set of ROI creation methods than any of the other tested toolboxes that were not specifically designed for this purpose alone. Therefore, it is important to recognize that fMROI is primarily dedicated to ROI creation and aims to integrate several desirable features from other software into a user-friendly interface.

A key distinguishing feature of fMROI is its ability to generate ROIs through an interactive graphical interface that includes an integrated viewer where users can visualize what they are designing along the creation process. This feature not only saves hours of work but also allows users to spot inconsistencies before storing the ROIs in NIfTI files. Furthermore, the design-time preview assists in creating more accurate ROIs. While MRIcroGL and FSL (via FSLeyes) offer a powerful viewer for inspection during ROI design, it offers limited options for ROI algorithms (spherical ROIs, voxel-intensity-based ROIs, and hand-drawn ROIs). In contrast, AFNI provides several algorithms but only the manual drawing tool is supported by GUI. SPM offers a simple GUI for parameter entry but lacks visualization at design time.

Although the fMROI algorithms were primarily developed for use within the GUI, they are not limited to it, as all ROI creation functions are accessible via the command line. This flexibility allows users to create scripts and automate routines, which can be especially beneficial for large-scale processing, such as applying a method that creates an ascending *regiongrowing* ROI using the voxel with the highest value in the right fusiform gyrus as a seed for multiple subjects. While this routine can be executed within the GUI, scripting proves to be a more efficient option in cases with a large number of subjects.

fMROI has also demonstrated its efficiency in combining ROIs using logical operations, as evidenced by qualitative analysis in figure 3C. In routine work, it is common to search for overlapping ROIs, for example, brain activations evoked by different tasks or conditions, or even brain activations across subjects (Ethofer et al., 2012; Fedorenko et al., 2010; Gerchen & Kirsch, 2017; Nichols et al., 2005; van der Zwaag et al., 2009). fMROI allows for performing logical operations directly from the GUI and also offers the option to run them through the command line. An interesting feature is the ability to combine not only images but also ROIs that are still under construction, further enhancing the software’s versatility.

One limitation of the current version of fMROI (1.0.0) is that it may experience a slow-down in performance when loading multiple images simultaneously, resulting in delays when updating the images and clicking buttons. This limitation may arise from the use of the SPM12 toolbox, which, although extremely robust and stable, is not optimized for real-time operations involving multiple images. However, it is important to note that fMROI was primarily developed for creating ROIs, which typically involve only a few template images. Consequently, this limitation does not significantly impact the broader functionality of ROI creation. Nevertheless, improvements to the handling of multiple images simultaneously is a priority for upcoming versions. We plan to develop new image manipulation and windowing strategies to enhance the viewer’s performance. Additionally, we aim to test and incorporate recent Matlab toolboxes, such as Medical Imaging and App Designer, if they prove stable and provide added functionality to the project.

Overall, fMROI presents a user-friendly software solution for ROI creation and brain image visualization. It offers several advantages, demonstrates comparable performance to established neuroimaging software, and provides a platform for collaborative work. Future updates and improvements will focus on enhancing the capabilities of the viewer and incorporating relevant Matlab toolboxes to further expand the functionality of fMROI.

## 6. CONCLUSION

In conclusion, here we present the development and validation of fMROI, a user-friendly software tool for creating ROIs and visualizing brain images. The software, implemented in Matlab, offers an intuitive GUI for complex ROI creation and incorporates functions for integrating user-specified algorithms. Comparative analysis with established neuroimaging software demonstrated fMROI’s comparable performance. Noteworthy features include the interactive graphical interface with a built-in vewer for real-time ROI design, support for command-line usage, efficient combination of ROIs through logical operations, and its collaborative platform design. Overall, fMROI provides a valuable tool for researchers in the field, enabling efficient ROI creation and facilitating neuroimage analysis.

## DATA AND CODE AVAILABILITY

For fMROI code access, visit https://github.com/peresasc/fmroi, and for a comprehensive user guide, please follow this link: https://fmroi.readthedocs.io.

To access the codes and datasets used to test the performance of the ROI creation algorithms, visit the Zenodo repository (https://zenodo.org/doi/10.5281/zenodo.10845324), or directly on GitHub (https://github.com/peresasc/fmroi_supplementary_data).

## AUTHOR CONTRIBUTIONS

**AP (corresponding author):** Conceived the project, developed the software, created the user guide webpage, conducted all validation analyses, created the scripts for ROI creation in AFNI software, and wrote the manuscript (original draft and revisions). **DV:** Assisted in conceptualization, validation analyses, programmed the codes for ROI creation in SPM software, conducted experiments with manual ROI drawing algorithms, and wrote the manuscript (revisions). **IV:** Assisted in creating the user guide webpage, assisted in validation analyses, programmed the codes for ROI creation in FSL software, and wrote the manuscript (revisions). **MM:** Assisted in creating the user guide webpage, assisted in validation analyses, created all ROIs in MRIcroGL software, and wrote the manuscript (revisions). **JW**: Assisted in conceptualization, validation analyses, and wrote the manuscript (revisions). **JA:** Assisted in project conceptualization, validation analyses, wrote the manuscript (revisions), and contributed to project supervision and administration.

## FUNDING

This work was supported by the European Research Council (ERC) under the European Union’s Horizon 2020 research and innovation programme Starting Grant number 802553; “ContentMAP’’ awarded to JA; by an European Research Executive Agency Widening programme under the European Union’s Horizon Europe Grant 101087584 “CogBooster” awarded to JA; by the Foundation for Science and Technology (FCT) under the strategic project of CINEICC with the reference UIDP/00730/2020, awarded to AP (call for proposals IT057-20-10242).

## DECLARATION OF COMPETING INTEREST

The authors have no conflicts to report.

## Notes

https://github.com/peresasc/fmroi

https://fmroi.readthedocs.io

https://zenodo.org/doi/10.5281/zenodo.10845324

https://github.com/peresasc/fmroi_supplementary_data

### Competing Interest Statement

The authors have declared no competing interest.

