## Supplementary material for "fMROI: a simple and adaptable toolbox for easy region-of-interest creation"

#### 1. SUPPLEMENTARY METHODS

##### 1.1 Generic GUI builder (autogen_gui.m)

In the absence of the GUI builder pair (i.e., the GUI and Caller functions), the fMROI uses a generic GUI builder (autogen_gui.m). In this case, each input argument from the method function generates an editable text box where the users can insert the input values. There are only four input argument exceptions because the fMROI uses special words to convey internal information to the ROI method functions. The keywords are: “srcvol” - the three-dimensional matrix containing the data of the image selected in the control panel; “curpos” - a three-dimensional vector containing the cursor's current position in data matrix coordinates; “minthrs” - value indicated in the minimum threshold slider in the image control panel; and “maxthrs” - value indicated in the maximum threshold slider in the image control panel.

When these special words are used as input arguments of the ROI method functions, the generic GUI builder does not create a text box, instead it directly inputs the predefined internal values defined by the fMROI GUI. This strategy simplifies conveying GUI information to new ROI creation algorithms, freeing developer collaborators from knowing GUI programming.

##### 1.2 Synthetic data creation:

To generate the set of three spheres, we first defined an equilateral triangle centered in the MNI coordinate [0,0,0] and corners [0,0,32], [28,0,-16], and [-28,0,-16] (data matrix in LAS [46,64,53], [32,64,29], and [60,64,29]) which results in edges of 56 mm. Then, we created three spheres centered in the triangle corners with radius equal to the triangle altitude (48 mm) and saved each sphere in an independent NIfTI file.

To generate the image with 64 spheres uniformly distributed, we defined a grid from the position [13, 13, 13] to [79, 79, 79] with lines 44 mm apart in x, y, and z coordinates. We centered one sphere at every line crossing with the radius defined randomly in the interval of 1 to 10 ensuring that the spheres did not touch each other. All elements of a sphere received an integer from 1 to 64, thus all voxel values of the first sphere are equal to 1, all voxel values of the second sphere are equal to 2, and so on.

The synthetic data with different shapes (here referred to as complex-shapes data) combines complex geometric shapes (not biologically inspired) with left and right hippocampi extracted from aparc+aseg atlas. In detail, it is a NIfTI file with 2 x 2 x 2 $mm^3$ isovoxel composed of four basic shapes: left and right hippocampi, a tetrahedron, and a cone (see figure 1B-3 ). Therefore, this dataset is composed of geometries that, in theory, are challenging for the ROI algorithms (sharp corners, straight lines, spirals, etc.) and facilitate the interpretation of the ROI algorithms outputs.

All voxels in the left hippocampus have a value of 1 (yellow). The voxels in the right hippocampus have random values from 3 to 3.9 (translucent red) plus 50 other voxels linearly distributed from 3.91 to 4 (blue dots). The tetrahedron has edges of approximately 4 mm (20 voxels) with vertices placed in [47, 52, 65]; [47, 70, 65]; [38, 61, 47]; [56, 61, 47]. The voxels in the central portion have a value of 5 and its four corners have respectively 5.2, 5.4, 5.6, and 5.8. The cone has an 8 mm base (40 voxels in diameter) and 6 mm height (30 voxels) filled with random values from 9 to 10. In addition, it has two spirals and a straight line connecting them. The external spiral has values between 11 and 12 with higher values in the tip [46, 84, 50] linearly decreasing in the direction of the base. The internal spiral is inverted in relation to the external and has values from 7.17 to 8 linearly increasing from its tip [46, 84, 30] to its base. The line that links the two spiral tips goes from the internal spiral [46, 84, 31] to the external spiral [46, 84, 49] linearly decreasing from 7.16 to 7.

To generate the DMN (Default Mode Network) image with Gaussian noise, we first fixed the noise amplitude in 5dB, i.e., the average of the absolute values of non-zero voxel values divided by the square root of 5. We then generated random values with a Gaussian distribution centered at zero and a standard deviation approximately equal to the noise amplitude and added these values to each of the elements in the original image default_mode_association-test_z_FDR_0.01.nii.gz.

### 2. SUPPLEMENTARY RESULTS

#### 2.1 Spheremask

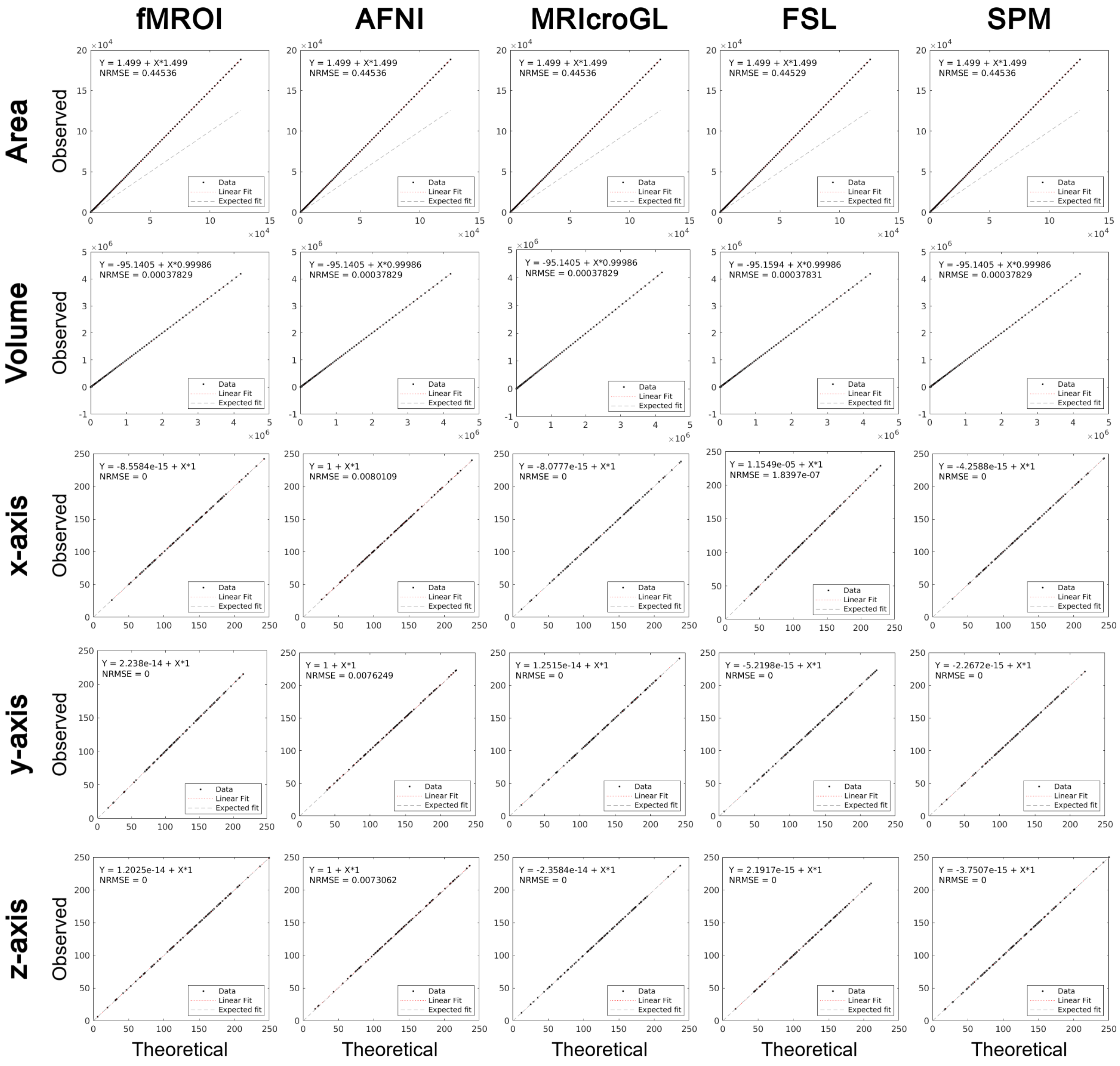


**Figure S1.** Linear regression of observed spherical ROI features against the respective theoretical data. The rows represent the assessed features, Area, Volume, and the sphere's center coordinates (observed - ROI center of mass; theoretical - nominal value entered into the ROI creation algorithm). The columns display all the tested programs: fMROI, AFNI, MRIcroGL, FSL, and SPM. In the upper left corner of each plot, the linear fit equation and the associated NRMSE are presented. The black points represent the observed data, the red dashed line represents the linear fit, and the gray dashed line represents the expected line in the absence of error (NRMSE = 0).

### 2.2 Cubicmask

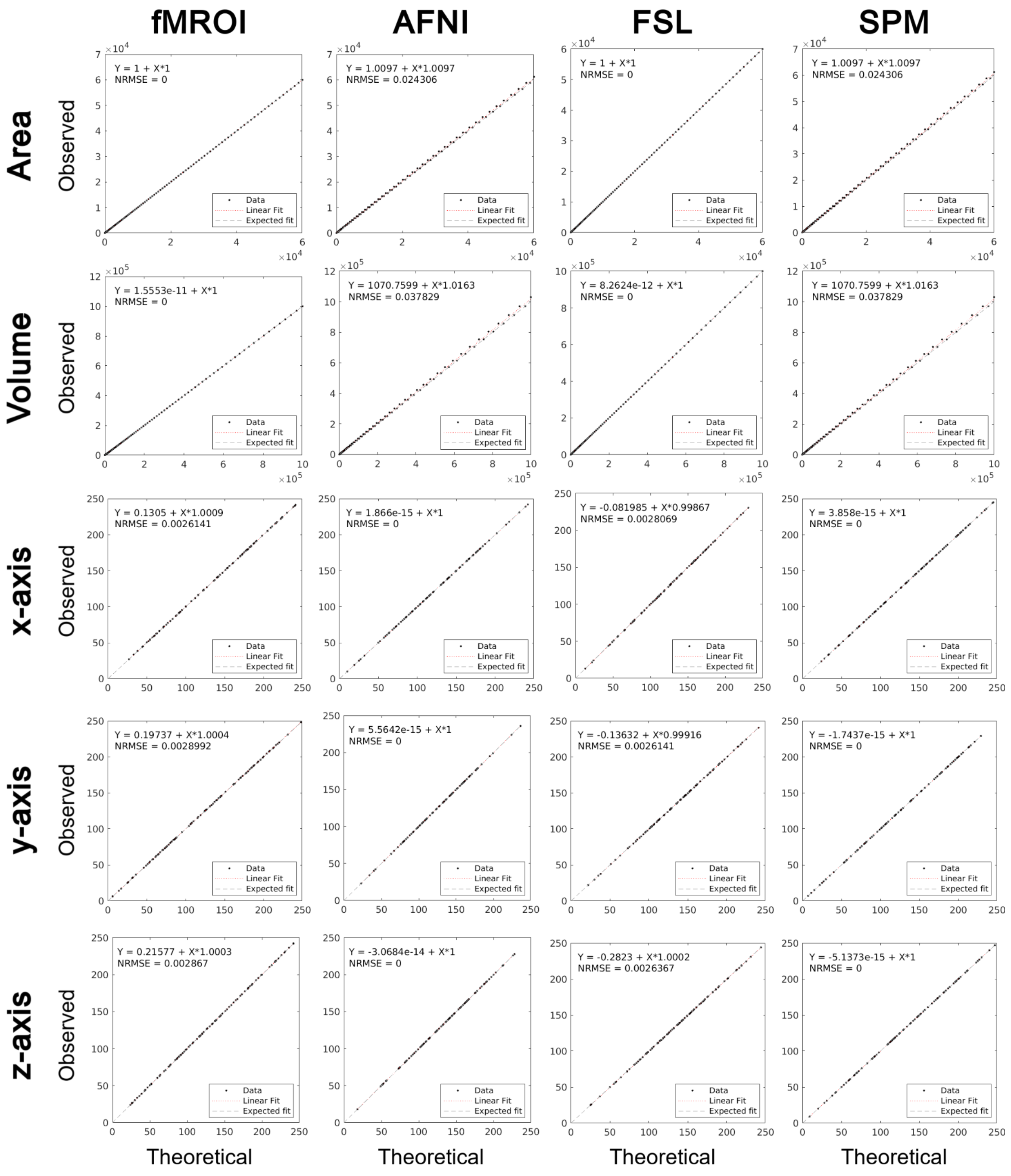


**Figure S2.** Linear regression of observed cubic ROI features against the respective theoretical data. The rows represent the assessed features, Area, Volume, and the cube's center coordinates (observed - ROI center of mass; theoretical - nominal value entered into the ROI creation algorithm). The columns display all the tested programs: fMROI, AFNI, FSL, and SPM. In the upper left corner of each plot, the linear fit equation and the associated NRMSE are presented. The black points represent the observed data, the red dashed line represents the linear fit, and the gray dashed line represents the expected line in the absence of error (NRMSE = 0).

#### 2.3 Cubicmask (only edges with odd number of voxels)

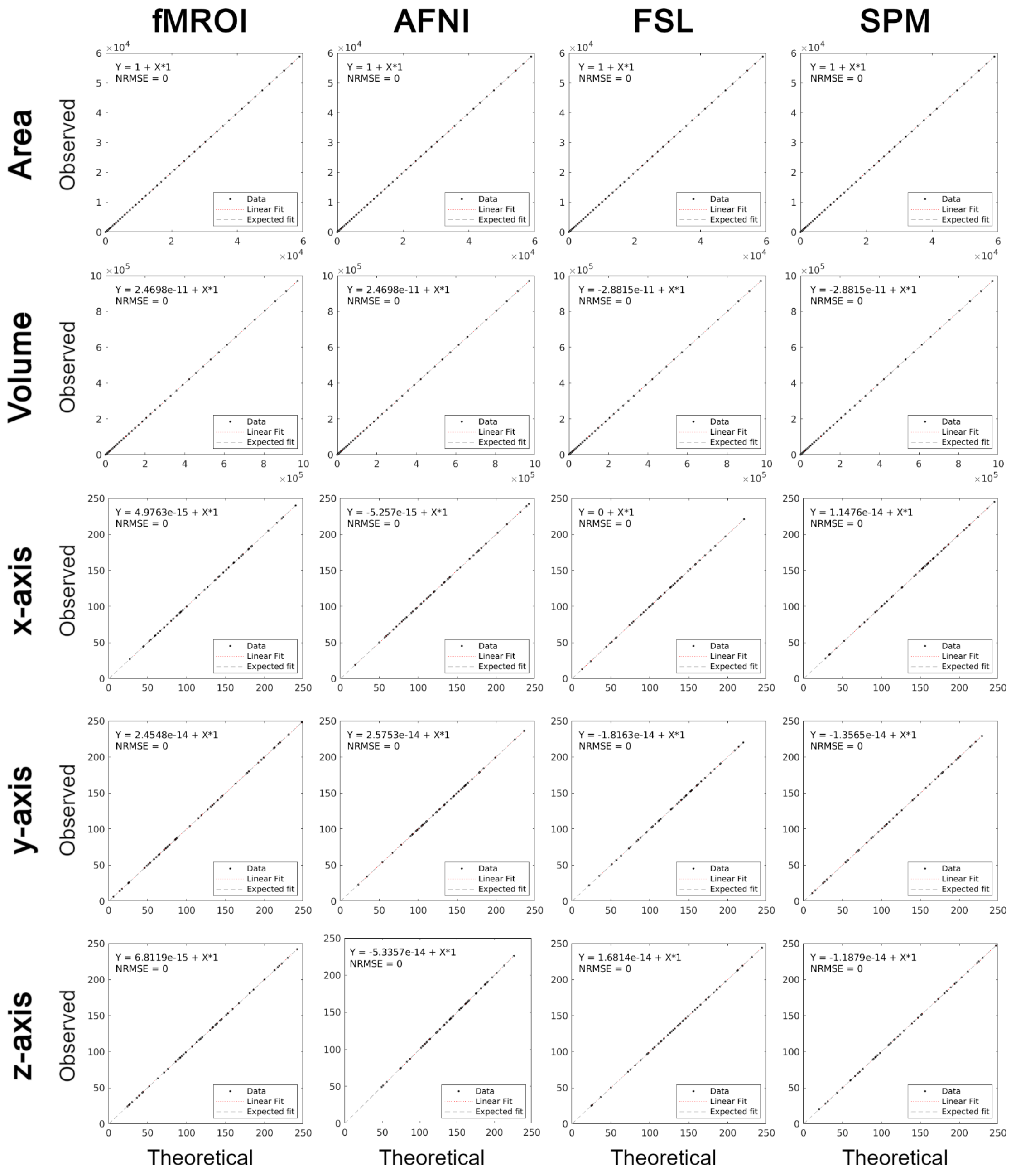


**Figure S3.** Linear regression of observed cubic ROI features (Limited to ROIs with an odd number of voxels on their edges) against the respective theoretical data. The rows represent the assessed features, Area, Volume, and the cube's center coordinates (observed - ROI center of mass; theoretical - nominal value entered into the ROI creation algorithm). The columns display all the tested programs: fMROI, AFNI, FSL, and SPM. In the upper left corner of each plot, the linear fit equation and the associated NRMSE are presented. The black points represent the observed data, the red dashed line represents the linear fit, and the gray dashed line represents the expected line in the absence of error (NRMSE = 0).
